# Towards a systematic map of the functional role of protein phosphorylation

**DOI:** 10.1101/872770

**Authors:** Cristina Viéitez, Bede P. Busby, David Ochoa, André Mateus, Marco Galardini, Areeb Jawed, Danish Memon, Clement M. Potel, Sibylle C. Vonesch, Chelsea Szu Tu, Mohammed Shahraz, Frank Stein, Lars M. Steinmetz, Mikhail M. Savitski, Athanasios Typas, Pedro Beltrao

**Author notes:** Contributed equally (Cristina Viéitez, Bede P. Busby). Correspondence to: Mikhail M. Savitski, Athanasios Typas and Pedro Beltrao.

## Abstract

Phosphorylation is a critical post-translational modification involved in the regulation of almost all cellular processes. However, less than 5% of thousands of recently discovered phosphorylation sites have a known function. Here, we devised a chemical genetic approach to study the functional relevance of phosphorylation in *S. cerevisiae*. We generated 474 phospho-deficient mutants that, along with the gene deletion library, were screened for fitness in 102 conditions. Of these, 42% exhibited growth phenotypes, suggesting these phosphosites are likely functional. We inferred their function based on the similarity of their growth profiles with that of gene deletions, and validated a subset by thermal proteome profiling and lipidomics. While some phosphomutants showed loss-of-function phenotypes, a higher fraction exhibited phenotypes not seen in the corresponding gene deletion suggestive of a gain-of-function effect. For phosphosites conserved in humans, the severity of the yeast phenotypes is indicative of their human functional relevance. This study provides a roadmap for functionally characterizing phosphorylation in a systematic manner.

## Introduction

Cells are constantly sensing and adapting to changes in environmental conditions. The relay of information from sensors to effector proteins often occurs via reversible protein post-translational modification (PTM), including protein phosphorylation. Regulation by phosphorylation allows cells to modulate protein activity, interactions and localization. In the past decade, mass spectrometry (MS)-based discovery of protein phosphorylation sites (phosphosites) has drastically changed our understanding of the extent of protein phosphorylation (Olsen *et al*, 2006). Recent estimates suggest that up to 75% of the yeast and human proteomes are phosphorylated, with over 10,000 and 150,000 non-redundant phosphosites described to date in the two organisms, respectively (Sadowski *et al*, 2013; Hornbeck *et al*, 2015). This vast extent of phosphorylation in eukaryotes has opened a debate on its physiological relevance (Landry *et al*, 2009; Lienhard, 2008; Kanshin *et al*, 2015; Beltrao *et al*, 2012), fueled by the fact that even for yeast we currently know the functional roles for less than 5% of these sites (Sadowski *et al*, 2013). It has been speculated that a significant fraction of phosphosites may have no function (Landry *et al*, 2009), consistent with the fast rate of evolutionary change in phosphorylation detected across species (Beltrao *et al*, 2009, 2012; Studer *et al*, 2016; Landry *et al*, 2009; Freschi *et al*, 2014). Evolutionary studies have suggested that between 35% to 65% of sites are constrained and therefore functional (Gray & Kumar, 2011; Landry *et al*, 2009). The degree of functional relevance and the functional role of most phosphosites, thus remain open questions.

Computational approaches have been developed to try to bridge the gap of functional assignment (Ochoa *et al*, 2019; Tunc-Ozdemir *et al*, 2017; Dewhurst *et al*, 2015). For example, prediction of phosphosites that may regulate protein interactions or cross-regulatory interactions with other PTMs have been tried (Nishi *et al*, 2011; Beltrao *et al*, 2012; Šoštarić *et al*, 2018). In addition, the conservation of phosphosites across species has been used to infer functionality, as phosphosites with well characterized functions are more likely to be conserved (Studer *et al*, 2016; Strumillo *et al*, 2019). In contrast, experimental approaches that functionally characterize PTM sites in a comprehensive manner are still lacking or are geared to a small number of phosphosites (Nakic *et al*, 2016; Oliveira *et al*, 2015).

Reverse genetics of gene deletion strains have been instrumental for the systematic elucidation of gene function (Tong *et al*, 2001; Costanzo *et al*, 2010; Collins *et al*, 2006). Similarly, reverse genetic screens of point mutants have shown promise for functional studies of protein regions (Braberg *et al*, 2013). Inspired by these studies, we set out to study the functional role of protein phosphorylation through a combination of chemical-genomics, molecular profiling and computational analysis. We created a yeast strain collection of phospho-deficient mutants that we screened for fitness phenotypes, together with the single-gene deletion library. In these assays 42% of the phospho-mutants displayed at least one significant growth phenotype when screened in 102 conditions, suggesting that these are likely functional. The correlation of the growth profiles across the 102 conditions, with those elicited by gene deletions was indicative of phosphosite function. We performed further characterization of the molecular changes elicited by some phospho-mutants by proteomics and lipidomics which further confirmed the inferences made from the genetic screen. Finally, the severity of the yeast phenotypes was indicative of the functional importance of orthologous human phosphosites.

## Results

### Construction and phenotypic profiling of a phospho-deficient mutant library

In order to study the functional role of protein phosphorylation using chemical-genomic approaches we generated a library of phospho-deficient mutant strains in *S. cerevisiae* **(Table S1)**. A compilation of experimentally determined *in vivo* phosphorylation sites was used to select 500 unique phosphosite positions including sites estimated to have different evolutionary ages (Studer *et al*, 2016). A total of 474 unique phospho-mutant strains, in 125 diverse genes, were generated with each position mutated to alanine at the genomic locus using a nearly scarless approach (Khmelinskii *et al*, 2011) (Fig. S1). To perform a quality control assessment of the generated phospho-mutant library we tested 50 strains for gene copy number by PCR and whole ORF sequencing: 5/49 (10%) showed additional PCR bands suggesting the potential acquisition of an extra gene copy and 5/42 (12%) showed additional non-synonymous mutations on the targeted ORF (Fig. S2). Overall the error rates associated with the library construction in this subset are relatively small. Erroneous strains are flagged in **Table S1** and were discarded, resulting in a library of 465 unique phospho-mutants.

The phospho-mutant library was arrayed as single colonies in a 1536 format and combined with the *S. cerevisiae* gene deletion library (4889 KOs) (Winzeler *et al*, 1999). These were plated on a panel of 102 different stress conditions that were selected to span a wide range of diverse perturbations (Fig. 1A and **Table S2**), including environmental, chemical or metabolic stresses (e.g. heat, DNA damage or nutrient starvation), drugs (e.g. antibiotics) and combination of stresses. Colony size was used as a proxy of fitness and a deviation from expected growth of each strain in each condition was computed using the S-score (Collins *et al*, 2006) (Methods, Fig. 1A). Positive and negative S-scores indicate that the mutant has increased or decreased resistance to stress, respectively. The resulting growth profiles (i.e. S-scores across all conditions) are shown as clustered heatmaps, separate for the phospho-mutants and KOs, indicating a diverse set of responses for both groups (Fig. 1B, available in **Table S3**). The number of mutant phenotypes per condition are shown in Fig. S2.

**Fig. 1.**
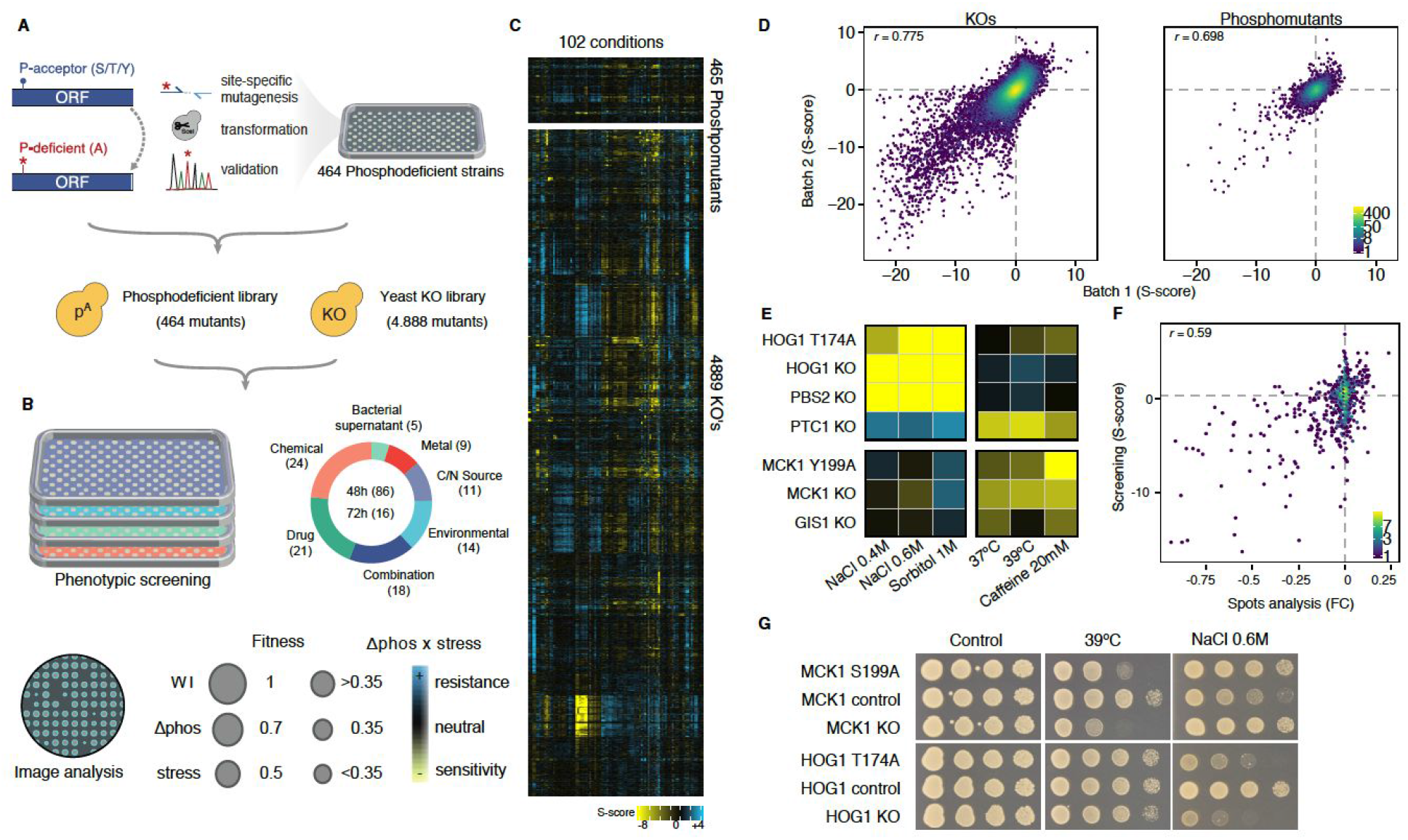
Phospho-mutant library construction, chemical genomic screen and quality control. **(A**) Phospho-mutant library construction (F**ig. S1 and Table S1**) and arraying of mutant libraries. (**B**) Chemical genomics screen against 102 conditions (**Table S2**), summary of image analysis and phenotype calling (See methods). (**C**) Overall heatmap of 465 phosphosites and 4889 KO’s in 102 conditions (**Fig. S3 and table S3)** (**D**) Correlation of S-scores for biological replicates of KO and phospho-mutants screened in 13 conditions (r Pearson correlation coefficient) (Table S2). (**E**) Heatmap of known regulatory phospho-mutants *HOG1-*T174A and *MCK1*-S199A, which phenocopy their corresponding knockouts. (**F**) Correlation between screen s-scores and fold changes from spot analysis for 762 pairs of mutant condition phenotypes (r=0.59; **Table S4).** (**G**) Example of serial dilution spot analysis, recapitulating the s-scores of *MCK1* and *HOG1* Control, KO and phospho-mutants in three conditions: untreated, heat and osmotic stress.

The measured growth profiles were highly reproducible as shown by the correlation of scores across biological replicates for gene deletions (Fig. 1C, r=0.775) and phospho-mutants (Fig. 1D, r=0.698). In addition, the measurements recapitulate expected mutant phenotypes. For example, deletions of either of two key kinases of the high osmolarity glycerol pathway (Hog1 and Pbs2) and a mutation of a key regulatory phosphosite in Hog1 displayed severe growth defects under osmotic shock conditions (Fig. 1E). In contrast, deleting the opposing phosphatase of the pathway (Ptc1) increased resistance in the same conditions. Similarly, we observed the expected sensitivity to growth under heat stress for deletion of Mck1 (Rayner *et al*, 2002) and a mutant of the regulatory phosphosite of this kinase (*MCK1* Y199A) (Fig. 1E and G). To further validate the high-throughput growth measurements, we performed independent, albeit less sensitive spot assays to test 762 pairs of mutant-condition phenotypes (Methods, Fig. 1F and **Table S5**). The spot assays were analyzed and given a semi-quantitative score that was largely consistent with the high-throughput growth phenotypes (Fig. 1F, r=0.59).

### Elucidating phosphosite function using growth profiles

Across all conditions, 42% of the phospho-mutants and 79% of the corresponding gene deletions showed at least one statistically significant phenotype (Fig. 2A). The number of phenotypes observed tended to be higher for phosphosites estimated to be older in evolutionary origin (Fig. 2A). Two observations suggest that more phosphosites may be functional than those captured by our assay: 6 out of 17 (35%) phosphosites with established functions did not show phenotypes (Fig. 2a); and rarefaction curve analysis indicated no saturation (Fig. S4), suggesting that additional conditions could yet reveal additional phenotypes. Given that the near scarless approach adds the 18 base pairs (bp) at the 3’-UTR region of each mutant, we asked whether this insertion could explain some of the observed phospho-mutant phenotypes. By measuring the average similarity of the mutant growth profiles with a control strain having only the 18 bp insertion (Fig. S5, **Table S6**) we identified 3 genes (*CHD1*, *RPB2* and *SWI4*) where the insertion may account for phenotypes. These 3 genes were discarded from further analyses but the fraction of phospho-mutants with phenotypes remains similar (40%).

**Fig. 2.**
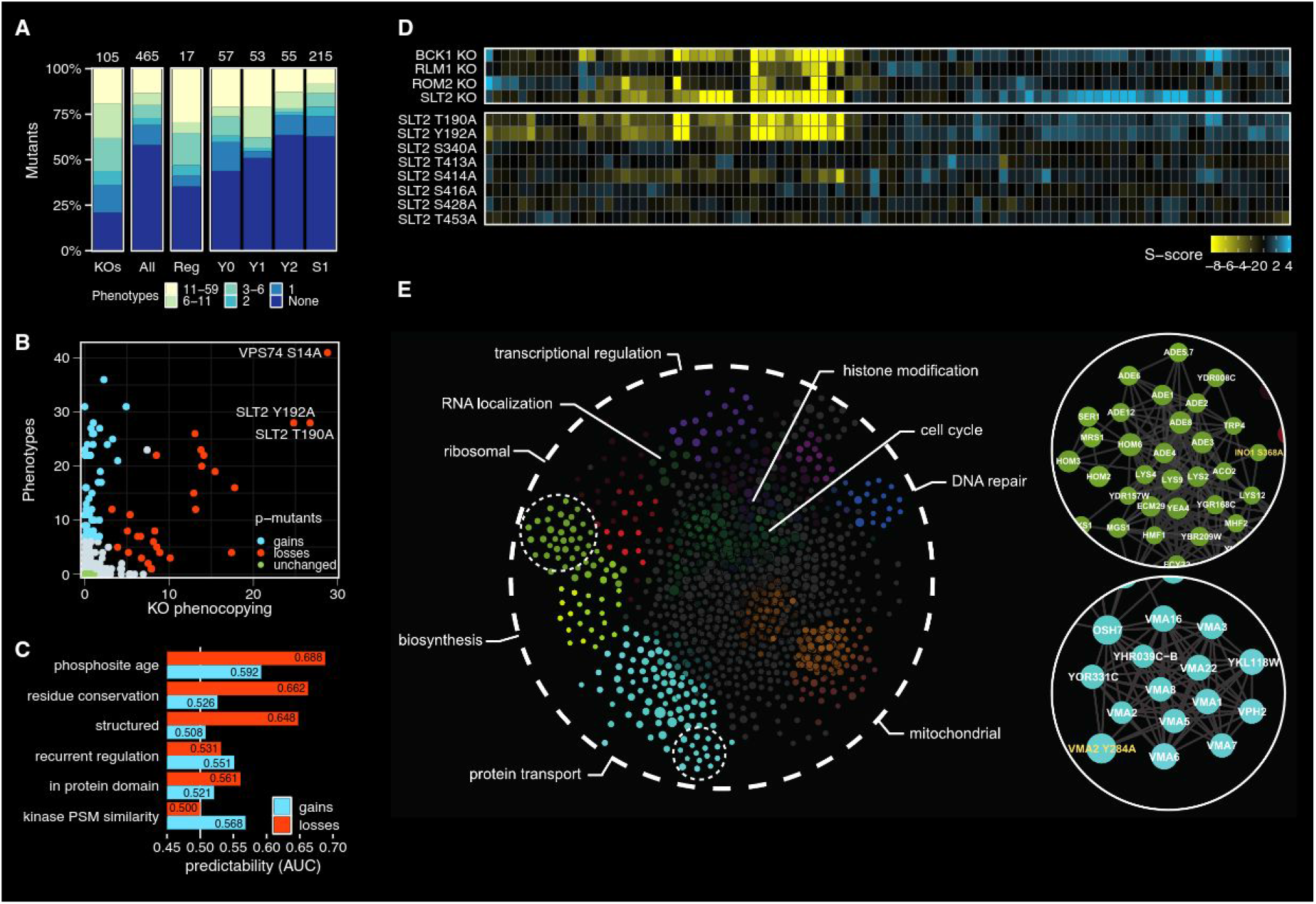
Chemical genomic screen functional analysis. (**A**) The percentage of mutants (KO or phospho-deficient) which have growth phenotypes (**Table S3**): KO: 105 non-essential available gene KO’s for which we constructed phospho-mutants; All: 465 phospho-mutants in our library; Reg:17 phospho-mutants previously described to be involved in protein function; Y0 to S1: phospho-mutants grouped based on their evolutionary age from more to less conserved: Y0 - 731 million years (My), Y1 - 434 My, Y2 - 61 to 182 My and S1 - <61 My (**B**) Growth fingerprints of *SLT2 KO* and eight phospho-mutants created in *SLT2.* Top: Heatmap depicting the phenotypic fingerprint of *SLT2 KO* which clusters with other related genes KO’s (*BCK1*, *RML1* and *ROM2).* Bottom: Unclustered phenotypic fingerprints of eight *SLT2* phospho-mutants. (**C**) Phospho-mutants that phenocopy their corresponding gene KO. Phenotypes - the number of significant phenotypes in the screen. KO phenocopying - log(*p*-value) of the correlation of the growth profiles between the phospho-mutant and corresponding gene-deletion. Blue = Gain of Function; red = Loss of Function; green = unchanged; grey=non-classified. (**D**) Area under the ROC curve (AUC) for the prediction of GoF or LoF classification, relative to unchanged, based on the following categories: phosphosite age (evolutionary conservation); residue conservation; position on protein structure; recurrent regulation; conservation of protein domain; or kinase recognition (PSM) site similarity. Blue = GoF, Red = LoF **(Table S6**). (**E**) SAFE analysis representation examining the functional organisation of phospho/gene KO network into functional gene ontology clusters. Highlighted in green and blue, are two examples which show significant enrichment in the categories of ribosomal biosynthesis (*INO1*-S368A) and protein transport (*VMA2*-Y284A).

Previous studies of gene deletion phenotypes have shown that the growth profiles across a large set of perturbations can be compared to assign functions to genes with unknown function based on a guilt-by-association principle (Beltrao *et al*, 2010; Tong *et al*, 2001; Costanzo *et al*, 2010; Collins *et al*, 2006; Nichols *et al*, 2011). The same principle can be applied to the study of single point mutants of phosphosites. For example, two point mutants in the Slt2 kinase (T190A and Y192A) show growth phenotypes that are highly similar to those elicited by the deletion of *SLT2* itself or the deletion of other members of the Slt2 cell wall integrity pathway (Fig. 2B). These two positions are known to be critical for the full activation of the Slt2 kinase (Martín *et al*, 2000). Another mutation (S414A) caused milder growth phenotypes in a subset of conditions, suggesting partial loss of function, whereas the remaining 5 point mutants did not show significant phenotypes. This result illustrates how the similarity of the growth profiles can be used for functional annotation of the phosphosites.

We classified each phospho-mutant according to the number of phenotypes and the extent by which the single point mutation shares the phenotypes (i.e. phenocopies) of the corresponding gene deletion (Fig. 2c). We identified a set of 29 phospho-mutants that phenocopy the KO (“loss-of-function” mutants) and an additional set of 55 mutants that interestingly have a large number of significant growth phenotypes that are not shared with the corresponding gene KO (“gain-of-function” mutants). These gain-of-function mutants have growth profiles that can in turn correlate with other mutants which can be suggestive of their function. For each phosphosite mutated in our screen, we collected annotations related to conservation, condition specific regulation and structural properties **(Table S7)**. We then asked if any of these features are characteristic of either the gain or loss of function group relative to a set of phosphosites without any significant phenotypes (Fig. 2D). Both functional groups showed increased conservation relative to the no-phenotype group but the loss-of-function group was particularly more likely to have more conserved phosphosites. Relative to the no-phenotype group, the loss-of-function phosphosites show enrichment for being within ordered regions, whereas the gain-of-function phosphosites are more likely to show a better match to known kinase specificity motifs.

In order to assign a putative functional role to each phosphosite, we computed all pairwise correlations of phenotypic profiles and built a network where two mutants are connected by an edge if they have significantly similar profiles (Fig. 2E). Based on this analysis, we provide a putative functional classification to phospho-mutants using the functional annotations of genes with similar growth profiles (**Table S8)**. As expected, the mutants tended to group in the network based on their functional characteristics (Fig. 2E) as identified by the Systematic Functional Annotation (SAFE) algorithm (Baryshnikova, 2016). In this network we find cases such as the loss-of-function *VMA2-Y284A* with phenotypes that resemble the deletion of *VMA2* itself or other members of the vacuolar H+-ATPase complex. In contrast, the *INO1-S368A* mutant has phenotypes that resemble that of the deletion of several amino-acid biosynthesis genes instead of the deletion of *INO1* itself, due to *INO1-S368A* having only partially overlapping phenotypes with the *INO1* deletion.

### Molecular changes relate to growth profiles and further elucidate phosphosite function

In order to associate the functional annotations derived from the growth profiles with molecular changes, we characterized 9 phospho-mutant strains, whose growth phenotypes were validated using two independently generated clones. We used a combination of an in-depth proteomics methodology named two-dimensional thermal proteome profiling (2D-TPP) (Becher *et al*, 2016; Savitski *et al*, 2018, 2014), and lipidomics (Fig. 3A). In addition to the molecular data collected here (**Table S9**), we analyzed the TPP data collected previously on two of the mutants also included in this study (*TDH3-S149A* and *T151A*) (Ochoa *et al*, 2019). The proteomics measurements allowed us to compare the thermal stability and abundance of thousands of proteins of each mutant strain relative to WT (**Methods**). For the lipidomics we quantified with LC-MS/MS the changes in 181 predicted lipid species (**Table S11** and **S12**), from which 19 were manually validated as described in **Table S13**. We determined and compared the number of significant phenotypes and molecular changes in each mutant (Fig. 3B) noting a trend whereby mutants with the highest number of phenotypes had the largest number of protein and/or lipid changes. In particular, *VMA2-Y284A* and *SIT4-S209A* showed many molecular changes in these assays.

**Fig. 3.**
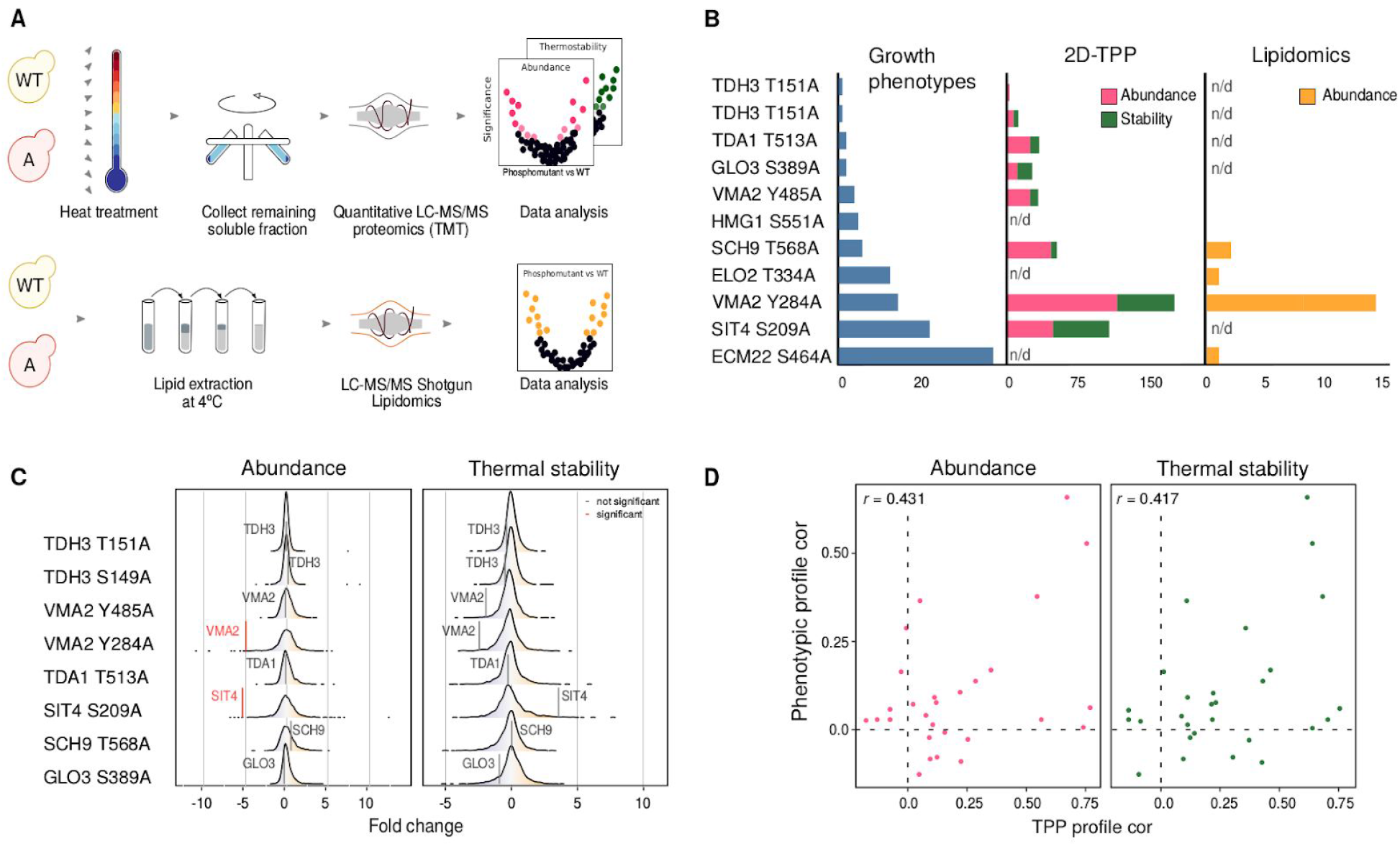
Molecular characterization of phospho-deficient mutants. (**A**) Graphical representation of the sample preparation, LC-MS/MS and data analysis for 2D-TPP (top) and shotgun lipidomics (bottom). (**B**) Summary of number of phenotypes observed in the genetic screen, significant hits in 2D-TPP and lipidomics per phospho-mutant. (**C**) Changes in protein abundance (left) and protein thermal stability (right) resulting from the phospho-mutation in their respective proteins. (**D**) Pairwise correlations of the phenotypic profiles of the phospho-mutants correlated with pairwise correlations in protein abundance (left) or thermal stability (right) generated by 2D-TPP.

The in-depth characterization of protein thermal stability allowed us to ask if the phospho-mutant proteins tend to change their thermal stability due to the alanine mutation. None of them showed a significant effect in thermal stability relative to WT, however *SIT4-S209A* showed a measurable increase in thermal stability that did not pass the significance cut-off (Fig. 3C). We next compared the distribution of changes in protein abundance and thermal stability of protein interaction partners of the mutated proteins and overall did not detect a significant difference with the exception of *VMA2-Y284A* and *SIT4-S209A* (Fig. S6), which were selected for further characterization. We then asked if the similarity of the phenotypic profiles for a pair of mutants is also reflected in similar profile of molecular changes. We observe that this is true when looking at the protein abundance (Fig. 3D, r=0.43, p-value 0.022) or thermal stability changes (Fig. 3D, r=0.41, p-value 0.027) but not apparent when looking at the changes in lipid composition (not shown). The lack of correlation with the lipid measurements may be due to fewer molecular measurements and fewer strains analyzed when compared with the proteomics experiments. These results suggest that at least for the proteomics measurements there is an agreement between the phenotypic and molecular characterization that should allow for a more in-depth mechanistic explanation of the mutational effects.

### Molecular characterization of phosphosite mutants in the vacuolar H^+^-ATPase Vma2 and phosphatase Sit4

Vma2 is a subunit of the vacuolar H^+^-ATPase (V-ATPase), a highly conserved proton pump critical for organelle pH homeostasis (Kane, 2006). For Vma2 12 phospho-mutants were screened with 6 having phenotypes. One of these (Y284) resulted in phenotypes similar to the deletion of Vma2, and several complex members (Fig. 4A) representing an example of a loss-of-function mutant. A second mutant of *VMA2* (Y485A), with a phenotypic profile that did not resemble the *VMA2* KO (r=0.70) was also selected for molecular characterization. Several growth phenotypes were validated by spot assays for these two mutants (Fig. S7).

**Fig. 4.**
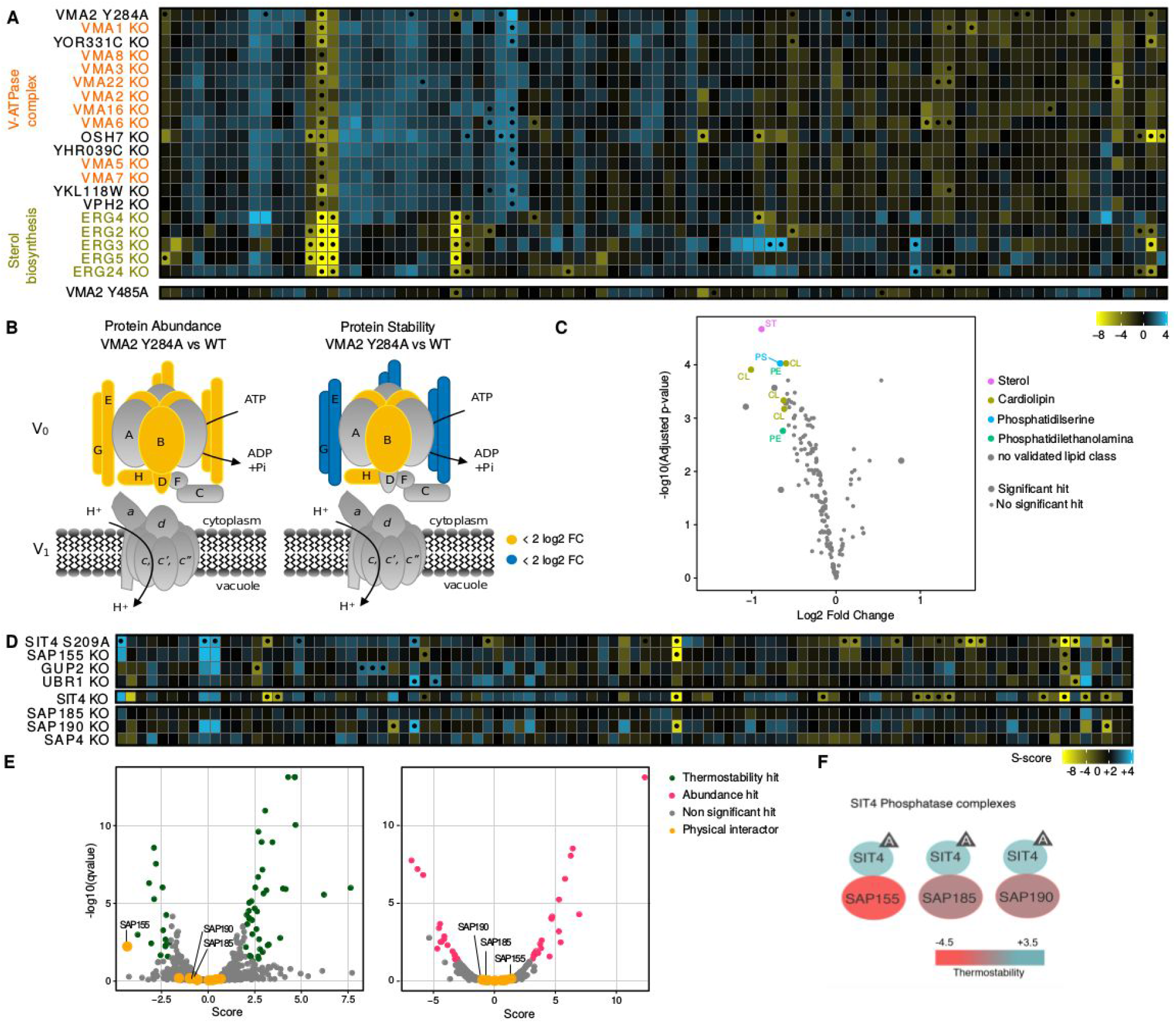
Molecular characterisation of Vma2 and Sit4 phospho-deficient mutants. (**A**) Top, growth profile of *VMA2-Y284A* which phenocopies *VMA2 KO* and clusters with members of the V-ATPase complex and members of the ergosterol biosynthesis pathway. Bottom, heat map (unclustered) of *VMA2-Y485A* which does not phenocopy *VMA2 KO.* (**B**) Graphical representation of the cytoplasmic V_0_ and membrane bound V_1_ subunits, colours represent complex members that significantly increase (blue) or decrease (yellow) in either protein abundance or protein stability measured by 2D-TPP **(Table S9**). (**C**) Lipid composition of *VMA2-Y284A* measured by shotgun lipidomics, values are Log_2_(fold change) compared to a control strain **(Tables S11, S12 & S13)**. Validated lipids **(Table S13)** in the following classes showed significant decreases: sterol (ST), cardiolipin (CL), phosphatidylserine (PS), phosphatidylethanolamine (PE). (**D**) Top, growth profile of *SIT4-S209A* in the same cluster with the deletion of *SAP155 KO*. Bottom, unclustered growth profiles of *SIT4 KO* and remaining phosphatase complex members*: SAP4, SAP190* and *SAP185 KOs*. (**E**) Volcano plots of *SIT-S209A* TPP data (**Table S9**). Score is the log2FC. Phosphatase complex members are labelled and physical interaction partners are shown in yellow. Significant changes in protein thermal stability (green) and significant changes in protein abundance (red) are shown. (**F**) Graphical Summary of the phosphatase complexes that Sit4 forms with SAP155 (destabilized), SAP185 (non-destabilized) and SAP190 (non-destabilized), colors are highlighting changes in protein thermal stability.

In the proteomics data the Y284A mutant showed a strong reduction in protein levels of Vma2 itself and in 5 out of 8 subunits of the V_0_ module (Fig. 4B), providing molecular evidence of the loss-of-function nature of this mutant. In contrast, Y485A showed no proteomic changes in *VMA2* or V-ATPase complex members, but it did show an increase in abundance of proteins enriched with annotations for mitochondrial localization (9/20) and transmembrane transport (GO:0055085, q-value<0.05; **Table S10**). In contrast, the Y284A mutant showed a decrease in abundance for proteins enriched in the same processes (GO:0055085,q-value<0.05; **Table S10**), suggesting different functional roles for these two phosphosites in Vma2 (Fig. S8). In the lipidomics data *VMA2-Y284A* showed a significant reduction in the abundance of 12 lipids (Fig. 4C) including zymosterol, a precursor of ergosterol, previously described to be necessary for V-ATPase activity (Vasanthakumar *et al*, 2019; Zhang *et al*, 2010). The reduction in zymosterol is consistent with the chemical genetic results, where we observed a high correlation between the *VMA2 KO*, *VMA2 Y284A* and 5 out of the 6 non-essential genes of the ergosterol biosynthesis pathway (Fig. 4A). In addition, the *VMA2-Y284A* mutant exhibited altered levels in 5 cardiolipin species, which have been previously suggested to play an important role in the maintenance, lubrication and/or activity the V-ATPase (Duncan *et al*, 2016; Zhou *et al*, 2011). These changes were not shared by the *VMA2* KO (**Table S14**) despite the fact that the KO and Y284A mutant shared most of the same growth phenotypes (Fig. 4A). Overall, the molecular data further confirms the LoF effect of the Y284A mutant and the difference in growth phenotypes elicited by the two VMA2 phospho-mutants.

Sit4 is a highly conserved PP2A-like phosphatase. Its activity and substrate specificity are regulated by the interaction with Sit4-associated proteins (SAPs) *SAP155*, *SAP185*, *SAP190* and *SAP4*. Deletion of all four SAPs together is equivalent to the deletion of Sit4, in terms of slower growth and cell cycle progression (reviewed in (Ariño *et al*, 2019)). One phosphosite position in Sit4 (S209) has been reported (Breitkreutz *et al*, 2010) and mutated in our study. The S209A mutant shows 23 growth phenotypes with a growth profile that correlates strongly with only one of the SAP KOs (SAP155, r=0.72) (Fig. 4D). While, S290A does not fully phenocopy *SIT4 KO*, it does share some of the same phenotypes such as reduced growth in heat stress and increased growth upon caffeine treatment. We also observed conditions where the S209A mutant had milder phenotypes than the *SIT4* KO such as reduced growth upon treatment with lipid biosynthesis inhibitors (ketoconazole and simvastatin). Several of these phenotypes were validated by spot assays (Fig. S9). In the 2D-TPP experiment, we observed a significant fraction of mitochondrial proteins changing in thermal stability, which agrees with the recently reported role of Sit4 regulating ATP synthase activity and mitochondrial function (Pereira *et al*, 2018). Sap155 was the only SAP protein showing a significant decrease in thermal stability in the S209A mutant. Together with the correlation of phenotypic profiles, this data suggests that S209A mutant has an impact on the Sit4-SAP155 phosphatase complex formation.

### Phospho-deficient library informs on the functional relevance of human phosphosites

We aligned the phospho-mutant genes to human orthologs identifying 72 yeast positions that aligned with 119 orthologous human phosphosites (as in (Ochoa *et al*, 2019)), due to one-to-many gene orthology assignments. To query whether measuring the phenotypes of phospho-mutants in yeast could inform us on the functional relevance of human phosphosites, we used human information that we have recently integrated into a single score of phosphosite functional relevance (Ochoa *et al*, 2019). We observed that human phosphosites that are predicted to be less tolerant to mutations (SIFT p-value<0.05) or have a high predicted functional score are more likely to have extreme phenotypes when mutated in yeast (Fig. 5A). These results suggest that we can use the yeast fitness data to prioritize and annotate functionally important human phosphosites.

**Fig. 5.**
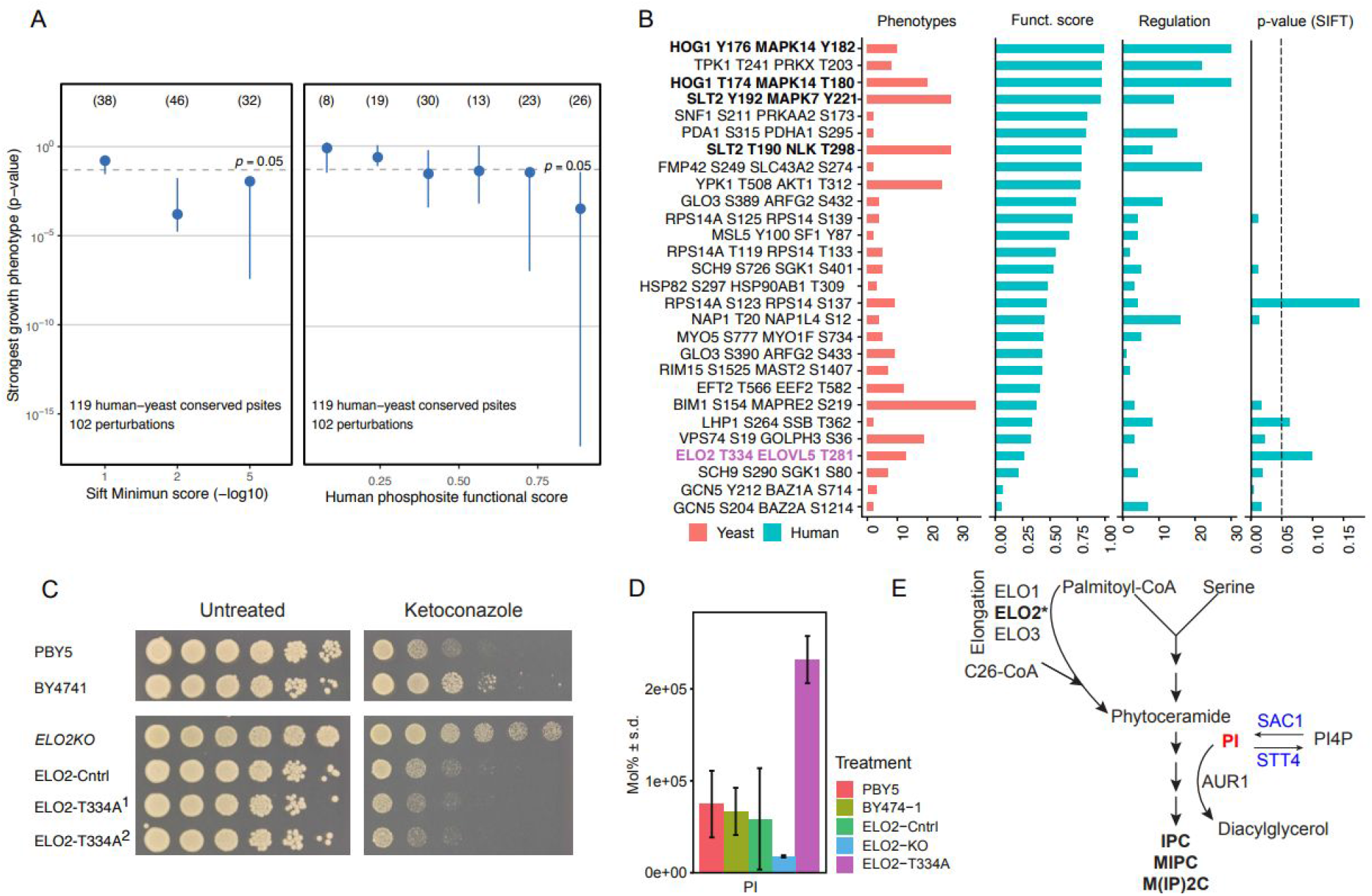
Yeast phospho-mutants with severe phenotypes can inform on the functional relevance of human phosphosites. (**A**) Human phosphosites orthologous to yeast phosphosites included in our screen were binned by SIFT and functional score and for each bin we represent the distribution of the impact of mutating the yeast phosphosites on yeast fitnest (p-value of the most significant phenotype for each phospho-mutant). (**B**) List of yeast (left) and conserved in human (right) phosphosites showing the number of yeast phenotypes in our screen and information on the human phosphosites. (**C**) Serial spot dilution assay for WT (PBY5 and BY4741), *ELO2 KO*, and two independent clones of ELO2-T334A phospho-mutant in Ketoconazole; (**D**) Molecular percentage of PI (18:1/26:1) in WT (PBY5 & BY4741), *ELO2*-control, *ELO2 KO* and *ELO2-T334A*; (**E**) Simplified schematic of sphingolipid biosynthesis pathway highlighting Elo2 (bold); the incorporation of PI (red); and known interactors Sac1 and Stt4 (blue).

From the 72 yeast phospho-deficient mutants that align to a human phosphosite, we selected those with at least 2 significant phenotypes. For one-to-many cases we selected the human position with the highest functional score (Fig. 5B). We obtained also annotations on the number of conditions in which the human sites are regulated (Ochoa *et al*, 2016), the predicted impact of mutations (Ng & Henikoff, 2003) and the phosphosite functional score (Fig. 5B **and Table S15**). This list contains human phosphosites with known functions such as the activation loop phosphosites of kinases (*MAPK14*-Y182/T180, *MAPK7*-Y221, *PRKX*-T203). To gain further insight into phosphosites conserved in human, two positions from this list (*GLO3*-S389 and *ELO2*-T334) were included in the molecular characterization (Fig. 3). We analyzed in more detail the T334A mutant of *ELO2* (a fatty acid elongase), which elicited a larger number of phenotypes (Fig. 5B) that were not strongly correlated with the corresponding gene deletion strain. A triple point mutant including this position had already been shown to impact on *ELO2* function (Zimmermann *et al*, 2013) but the role of T334 has not yet been investigated on its own. Some of the growth differences between *ELO2-T334A* and *ELO2* KO were validated by spot assays (Fig. 5C). For example, the gene deletion shows resistance to an inhibitor of sterol biosynthesis (ketoconazole) while the T334A shows a modest increase in sensitivity. We investigated the differences in lipid composition in these two mutants (**table S14**) finding a general trend of opposite impact on the phosphatidylinositol (PI) species, with an accumulation of PI (18:1/26:1) observed in the T334A and a depletion observed in the KO (Fig. 5D). Elo2 acts towards the start of the sphingolipid biosynthesis pathway, where it aids in the synthesis of very long chain fatty acids, upstream of where ceramide is combined with PI (Fig. 5E). Interestingly, *ELO2* deletion has interactions with the enzymes involved PI production and consumption - a negative genetic and physical interaction with *SAC1*, and is synthetic lethal with *STT4* (Cherry *et al*, 1997). These interactions are consistent with the observed perturbation in PI levels. The results indicate that the phospho-deficient mutant has an impact on the normal function of Elo2 and possibly acting in a gain-of-function fashion.

## Discussion

We have developed here a chemical-genetic approach that can be used to systematically identify phosphosites that strongly contribute to fitness in a condition specific manner. Conservation based studies have suggested that up to 65% of phosphosites may be non-functional (Landry *et al*, 2009). While, in our assay approximately 40% of the phospho-mutants displayed at least one significant phenotype, it is not straightforward to extrapolate this number to all phosphosites. The assays did not reach saturation as judged by the rarefaction curves or the lack of measured phenotypes for some phosphosites with known function. This could be due to a number of reasons - there could be additional relevant stress conditions that were not assayed; some phenotypes may only become relevant with phospho-mimetic mutations instead of alanine; and some phosphosites may work within clusters whereby one single mutation may be easily compensated by a nearby phosphosite. This would suggest that a larger fraction of phosphosites should be relevant for fitness. Nevertheless, we think this study may provide a lower bound estimate (i.e. ~40%) on the fraction of phosphosites that result in a growth phenotype when mutated. Importantly, in addition to finding which phospho-mutants have the strongest phenotypes, the correlation of the growth profile of the phospho-mutants with those elicited by the knock-out strains generated a rich source of functional annotations.

It is possible that some phenotypes are induced by destabilization of the protein by mutations that would not be physiological. However, all mutated positions were selected based on mass spectrometry evidence for *in vivo* phosphorylation. Such positions should be able to accomodate a phosphate group and therefore be less likely to be destabilized by an alanine mutation. In addition, we were able to measure thermal stability changes for 8 phospho-mutants with one showing an increase and one showing a decrease that were not statistically significant. Although this is a small sample size it suggests that the mutations themselves are not often causing destabilization of the proteins. Therefore the phenotypes measured should be, to a large extent, the result of the loss of function of that position (i.e. loss of phosphorylation).

Analysing different features of the phospho-mutants with phenotypes we observed that conservation was the best single predictor of functional relevance - in particular for the loss-of-function mutants - but conservation alone is not sufficient to identify most phosphosites with important phenotypes. This reinforces the need to develop machine learning based approaches to integrate different features (Dewhurst *et al*, 2015; Ochoa *et al*, 2019). Unbiased fitness measurements for a large number of phospho-mutants, as determined here, will be a crucial part of the further development of such predictors. While it is still not possible to perform large scale genome point mutant studies for human phosphosites there are several methods under development (Zafra *et al*, 2018) that may reach a point where they can be employed for large scale phosphosite studies. Despite the overall poor conservation of phosphorylation between human and yeast, these studies in yeast were shown here to be useful to annotate orthologous human positions. Finally, these approaches have been used here to study protein phosphorylation but they can be, in the future, be directed at other PTMs such as ubiquitination and acetylation, among others.

## Methods

### Strains and Media used

The S288C Mat**a** haploid KO library (4889 KO’s) and the Matα phospho-deficient mutant library (474 strains) were maintained on YPD+G418 and YPD respectively, prior to screening in 384 colony format. All other strains used for this study are displayed in **table S1** and were maintained on the appropriate selection media. Synthetic complete medium

### High Throughput yeast transformation

Yeast transformations were performed as previously described (Gietz & Schiestl, 2007) with some modifications. Briefly, transformations were carried out using chemical transformation in 96 well PCR plates and incubations were carried out in a thermocycler. Yeast cells were grown to exponential phase, harvested by centrifugation and washed with Lithium Acetate (Sigma)-TRIS (sigma) EDTA (sigma) buffer (LiAc 100 mM, 1x TE pH 8.0), resulting in chemically competent yeast cells. Up to 30 uL of DNA (either PCR product or plasmid) was added to each well followed by transformation mix (100 uL 50% PEG 3350) Sigma), 15 uL 1M LiAc, 20 uL single stranded carrier DNA from salmon sperm (Sigma)) and 30 uL competent cells. This mix was heat shocked at 42°C for 40 min and plated onto the appropriate selection medium, if selection is antibiotic resistance, an outgrowth of 4 hours at 30°C was carried out prior to selection.

### Phospho-deficient mutant library construction

Phosphodeficient point mutation (Serine/Threonine/Tyrosine to Alanine) were constructed using a modified homologous recombination method (Toulmay & Schneiter, 2006) in the strain Y8205 (*MAT*α *can1*Δ::*STE2*pr-Sp*HIS5 lyp1*Δ::*STE3*pr-*LEU2 LYS2 his3Δ1 leu2Δ0 MET15*+ *ura3Δ0*; a kind gift from Nevan Krogan; REF). Briefly, prior to introducing the specific point mutation, the *SCE1* endonuclease under the control of the galactose inducible promoter from plasmid pND32; (Khmelinskii *et al*, 2011)) was inserted at the *LEU2* loci. Next, the *URA3* selectable marker flanked by 18 bp recognition site (ATTACCCTGTTATCCCTA) for the SCE1 endonuclease was inserted after the stop codon in the 3’ UTR of 102 genes. insertion of the marker was confirmed by PCR. Next, the point mutation was introduced in a primer 55 bp in length upstream of the 3’UTR, this primer plus one in the centre of the URA3 marker was used to introduce the mutation and amplify the 5’ piece of DNA (Piece 1), additionally another PCR was carried out amplifying an overlapping piece of the URA marker with 55 bp homology to the 3’UTR (piece 2) as shown in Fig S1. Piece 1 and piece 2 were then simultaneously transformed into the WT Y8205 GALp-SceI strain. Point mutations were confirmed via sequence analysis of the point mutation containing region. The URA3 marker was removed via replica plating onto medium containing 2% galactose followed by a positive selection on 5-Fluoroorotic Acid (Boeke *et al*, 1987) to ensure the marker had been successfully removed **(see Fig.S1).** For quality control, 50 mutants were examined for gene copy number variation and additional mutations in the gene body (Fig. S2). All phosphodeficient mutants are listed in **Sup Table 1**

### Chemical genomics screen

Growth of the S288C (BY4741) K/O and phospho-deficient strain collection was evaluated on concentrations of chemical and environmental stress conditions (**Table S2**) that inhibit the growth of BY4741 by approximately 40%. The libraries were maintained and pinned with a Singer RoTor in 1536 colony format. Synthetic complete media (Kaiser et al 1994) was used with or without the stress condition, incubated at 30°C (unless temperature or medium was a stress eg Nati medium (CSM35), was prepared as described in (Ponomarova *et al*, 2017) for 48h or 72h, and imaged using a SPIMAGER (S&P Robotics equipped with a Canon Rebel T3i digital camera.

The screen was carried out in two main batches, consisting of 70 conditions in batch 1 and 29 conditions in batch 2, all conditions and concentrations are listed in **Table S2.** In order to account for variation of the method 13 conditions were tested in both batches (anaerobic growth, amphoteracin B, nystatin, DMSO, 2,4,D, glycerol, maltose, hepes buffered medium, caffeine, 6-AU, paraquat, 39°C, sorbitol).

### Chemical genomics data analysis

Raw plate images were cropped using ImageMagick to exclude the plate plastic borders; raw colony sizes were extracted from the cropped images using gitter v1.1.1 (Wagih & Parts, 2014), using the “autorotate” and “noise removal” features on. Poor quality plates were flagged when no colony size could be reported for more than 5% of colonies, which indicates poor overall quality, or when no colony size could be reported for more than 90% of a whole row or column, which indicates a potential grid misalignment. Known empty spots in each plate where used to flag incorrectly labelled plates. Conditions with less than three replicates across the two batches of the experiment were excluded from further processing. Raw colony sizes for the remaining conditions were used as an input for the S-score (Collins *et al*, 2006), with default parameters except the minimum colony size which was set to 5 pixels. The algorithm computes an S-score, which indicates whether the growth of each mutant/KO is deviating from the expected growth in each condition. The raw S-scores were further quantile normalized in each condition. Significant positive and negative phenotypes were highlighted by transforming the S-scores in Z-scores, given that the S-scores in each condition follow a normal distribution. P-values were derived using the survival function of the normal distribution and corrected using an FDR of 5% (false discovery rate). The whole dataset, of mutant-condition scores is available as **Table S3**.

The reproducibility of the chemical genomics screen was assessed by dividing the raw pictures according to the batch of origin; the EMAP algorithm was then used to compute a set of s-scores for each batch. For the 13 conditions that were tested in both batches the S-score correlation was computed for the phospho-mutants and the KOs separately. We refer to this analysis as both technical and biological replicate because the inoculates are derived from the same source plate but at very different times (Fig. 1D).

To associate phosphosites with specific functions we correlated the growth profile, defined as the S-scores across conditions, of each phospho-mutant with the profile of all gene deletion strains. We defined the significant phospho-mutant to gene correlations (positive correlation with 5% FDR cutoff) and performed an enrichment test for the top most enriched GO terms of these associated genes. We excluded GO terms that were too unspecific (i.e. annotating more than 500 genes or linked to 10 or more of phospho-mutants). To reduce redundancy in the annotations we also measured GO-term similarity by calculating Jaccard Similarity Coefficient between all the GO gene sets. For each phospho-mutant we then selected the top 2 associated GO terms more specifically associated with each mutant from different GO term similarity clusters. Specificity of a GO term was defined by the number of phosphosites associated with it with broader terms presumed to be associated with more phosphosites.

### Screen validation by serial dilution assay

Yeast were grown in 96 well microtiter plates in YPAD and incubated for 24 h until saturation. The strains were then serially diluted four times at a 1/20 dilution in fresh 96 well plates, the dilutions were performed using a liquid handler (Beckman Coulter BioMek FX^P^). The diluted cells were then immediately spotted onto stress condition agar plates using the VP405 96 (V&P Scientific) format manual pinning tool. The agar plates were incubated for 48 and 72 hours at 30ºC, unless other temperature is indicated. Agar plates were imaged using a SPIMAGER (S&P Robotics equipped with a Canon Rebel T3i digital camera). The resulting spot assays were scored 0-5 by eye based on colony sizes compared to controls on each plate.

### Two-dimensional thermal proteome profiling (2D-TPP)

The 2D-TPP protocol (Savitski *et al*, 2014) was adapted to be compatible with *S.cerevisiae*. Yeast cells were grown overnight at 30°C in YPAD and diluted to OD 0.1 in 50 ml fresh YPAD media. Cultures were collected by centrifugation at 4000x g for 5 min when OD_600_ reached ~0.7, immediately frozen in liquid nitrogen, washed with PBS and resuspended to an equivalent of OD_660_ of 125. 20ul was aliquoted to 10 wells of a PCR plate and the plate was centrifuged at 4.000g for 5 min and was subjected to a temperature gradient for 3 min in a thermocycler (Agilent SureCycler 8800) followed by 3 min at room temperature. Cells were lysed in 30 ul of cold lysis buffer (final concentration: Zymolyase (Amsbio) 0.5mg/ml, 1x protease inhibitor (Roche), 1x phosphatase inhibitor PhosSTOP (Roche), 250U/ml benzonase and 1mM MgCl2 in PBS for 30 min shaking at 30°C, followed by five freeze-thaw cycles (freezing in liquid nitrogen, followed by 30s at 25°C in a thermocycler and vortexing). Protein digestion, peptide labelling, Mass spectrometry-based proteomics and data analysis were performed as previously described (Mateus *et al*, 2018).

### Lipid Extraction

Yeast cultures were grown in 5 mL synthetic complete media to an OD_600_ of 1.0, harvested by centrifugation and washed with 1 mL dH_2_0, transferred to a 1.5 mL tube and the cells were pelleted by centrifugation. The following extraction process was carried out at 4 °C. Sterols were extracted from the cellular pellet by the addition of 150 µL acid washed glass beads along with 900 µL chloroform: methanol (1:2), vortexed vigorously for 10 mins and then incubated at 4 °C for 15 hours with shaking at 1100 rpm. After incubation a further 300 µL of chloroform and 300 µL of dH_2_0 was added, the mixture was vortexed for 30 sec, rested for 2 mins then centrifuged for 5 mins at 9,000 x g. The solvent fraction was removed and transferred into a new 1.5 mL tube. A second extraction was performed by the addition of 600 µL chloroform, vortexing vigorously for 30 sec followed by a final centrifugation for 5 mins at 9000 x g, the solvent fraction was removed and added to the previous solvent fraction. The combined solvent fractions were then dried over nitrogen and stored at −80 °C.

### Lipidomics LC-MS/MS Analysis

Lipid extracts were separated on a Kinetex C18 2.1 x 100 mm, 2.6 µm column (Phenomonex, Aschaffenburg, De). Separation was achieved by gradient elution on a binary solvent Vanquish UHPLC (Thermo Fisher Scientific, Bremen, DE). Mobile Phase A consisted of ACN: H2O (60:40) while mobile phase B consisted of IPA: ACN (90:10). For positive ionization, the mobile phases were modified with 10 mM ammonium formate and 0.1% formic acid while for the negative ionization mode, the mobile phases were modified with 5 mM ammonium acetate and 0.1% acetic acid. A flow rate of 260 µL/min was used for the separation and the column and sample tray were held constant at 30ºC and 4 ºC respectively.

### Lipidomics Mass Spectrometry Instrumentation

MS analysis was performed on a Q-Exactive plus Mass Spectrometer (Thermo Fisher Scientific, Bremen, DE) equipped with a heated electrospray ionization probe. In both the positive and negative ionization modes, the S-Lens RF level was set to 65, the capillary temperature was set to 320 °C, the sheath gas flow was set to 30 units and the auxiliary gas was set to 5 units. The spray voltage was set to 3.5 kV in the negative ionization mode and 4.5 kV in the positive ionization mode. In both modes, full scan mass spectra (Scan Range m/z 100 −1500, R=35K) were acquired along with data-dependent (DDA) MS/MS spectra of the five most abundant ions. DDA MS/MS spectra were acquired using normalized collision energies of 30, 40, and 50 units (R= 17.5K and an isolation width = 1 m/z). The instrument was controlled using Xcalibur version 4.0.

### Data analysis, lipid annotation and confirmation

Progenesis Q1, version 2.0 (Non-Linear Dynamics, A Waters Company, Newcastle upon Tyne, UK) was used for peak picking and for chromatographic alignment of all samples (with a pooled sample acquired in the middle of the LC-MS sequence used as a reference). Lipids were initially annotated from Progenesis Metascope and LipidBlast databases. Putative identification of lipids was done for ions that had MS/MS data. A pooled sample was used for the lipid identification of statistically significant lipids. An inclusion list was created to acquire MS/MS spectra of these lipids and spectra were acquired as described above. Full scan spectra of the pooled sample were also acquired by polarity switching between the positive and the negative ionization modes. Raw MS/MS spectra were log transformed and distribution of spectra were compared to remove any outlier samples. Differential analysis of the spectra were performed using Limma (Ritchie *et al*, 2015) for each phosphomutant or KO with their respective controls.

---

**Fig S1.**
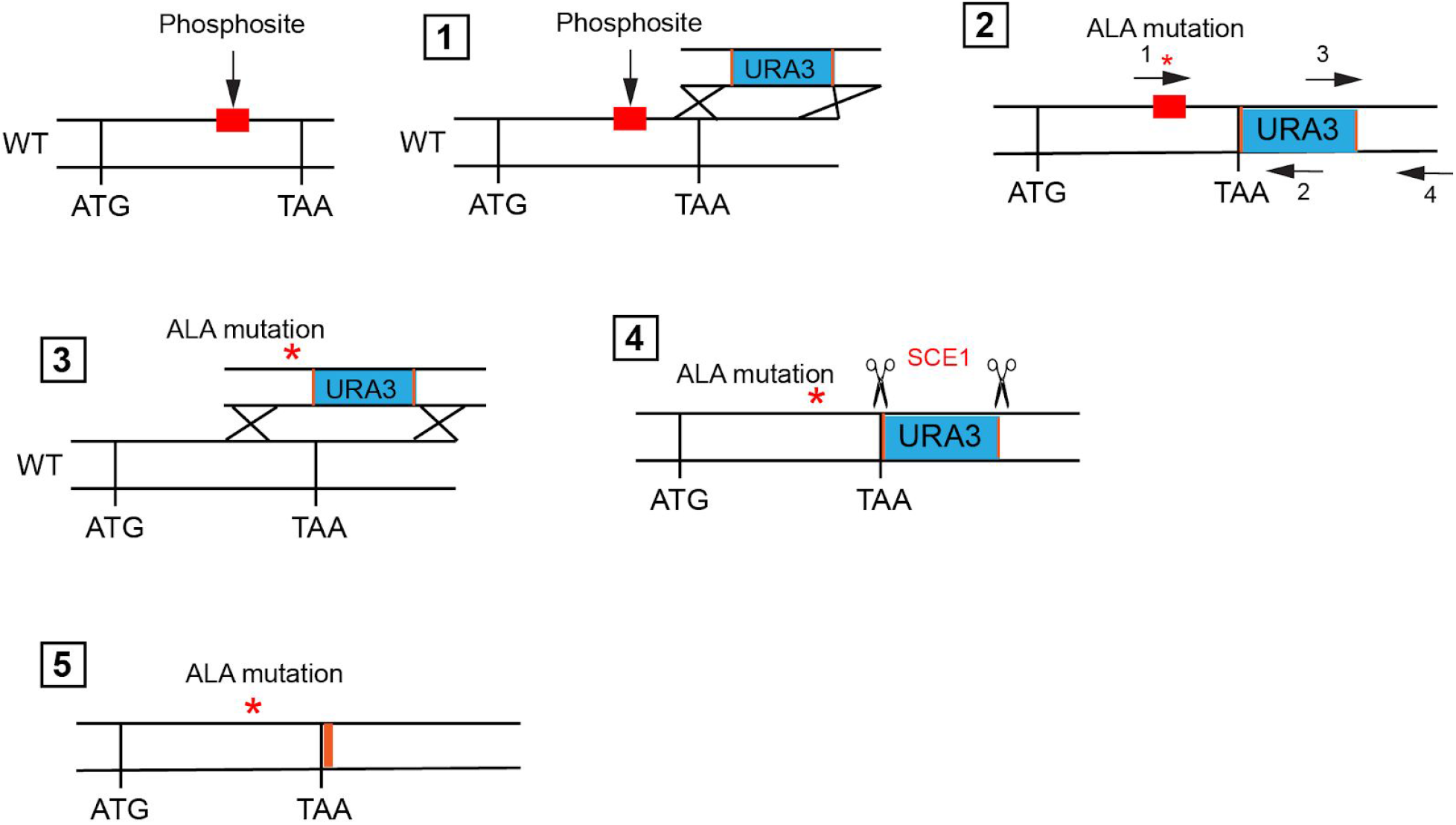
Phospho-mutant library construction. Schematic of how the point mutation was introduced and removal of the *URA3* marker: 1) Insertion of the *URA3* marker from pAG60 in the 3’UTR. 2) Amplification of the point mutation along with the 5’ end of the *URA3* marker using primer 1 (containing the point mutation) and 2 (*URA3* internal)in addition to the overlapping PCR product of the 3’ end using primers 3 and 4. 3) insertion of the point mutation in along with the *URA3* marker into a clean strain. 4) removal of the marker via activation of the SCEI endonuclease by plating onto galactose containing media. 5) The resulting markerless (18 bp SCEI recognition site remains) phospho-mutant.

**Fig S2.**
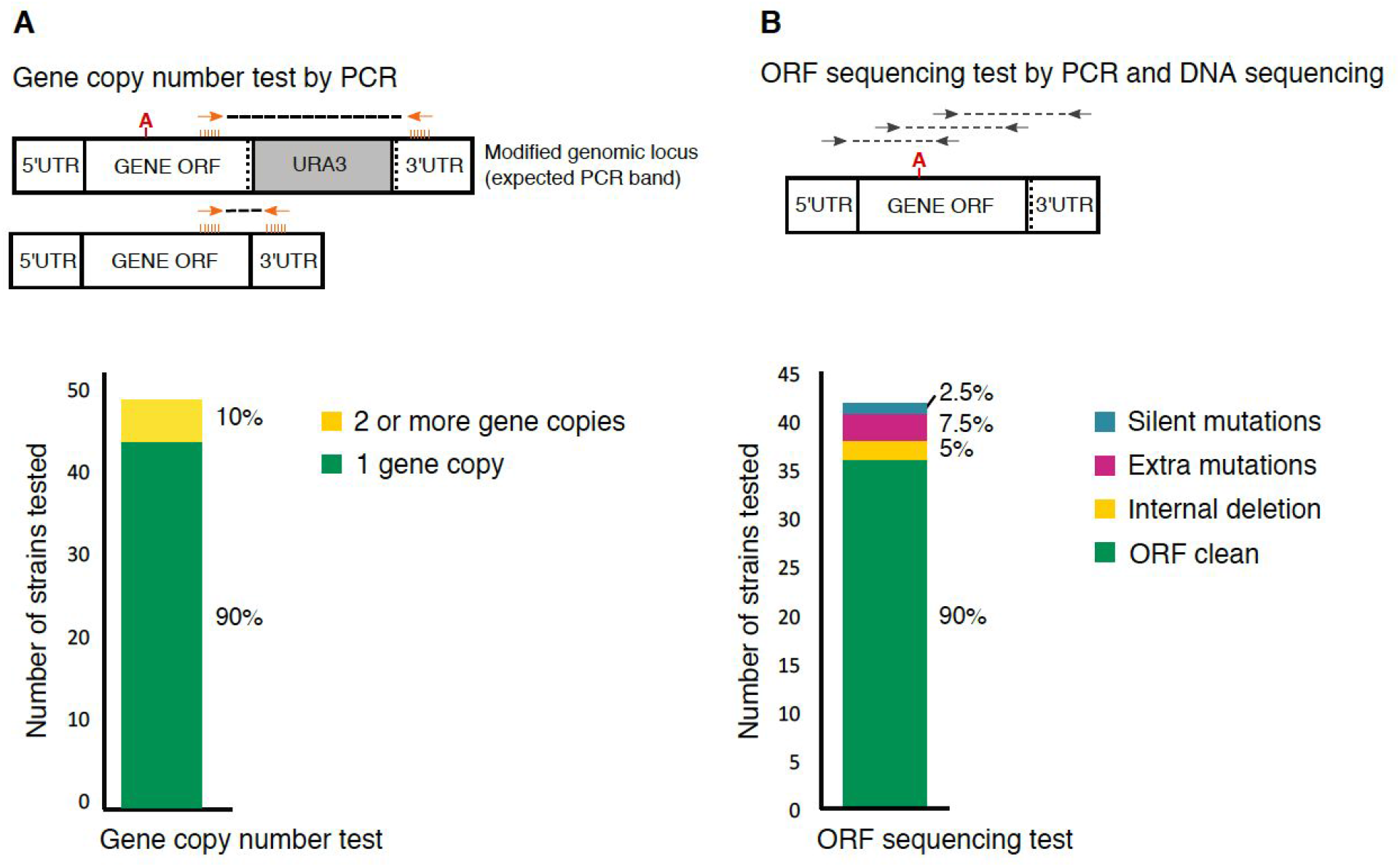
Quality control of the phospho-mutant library. A selected subset of 50 strains were checked by **A)** PCR on the strain that contains the URA3 marker (before removal, see Fig S1). We use PCR to amplify a fragment from the gene open reading frame (ORF) to the gene 3’UTR. The expected PCR size corresponds each gene specific 3’region and the addition of 1.4Kb corresponding to the URA3 marker cassette introduced after the stop codon. From the 49 strains for which we got PCR products: 44 show the expected PCR size (1 gene copy) and 5 show the expected PCR size and an additional band that was 1.4Kb smaller, corresponding to the gene in the absence of the URA3 cassette, suggesting that the gene got duplicated (2 or more gene copies). **B)** DNA sequencing of the ORF region. We observed a clean ORF containing only the desired mutation in 36 cases (90%), an internal deletion of few base pairs in 2 cases, an extra non-synonymous mutation in 3 cases and the presence of an additional synonymous mutations in 1 case. In 9 cases we did not get good enough quality of the sequence to discard the presence of an extra mutation.

**Fig S3.**
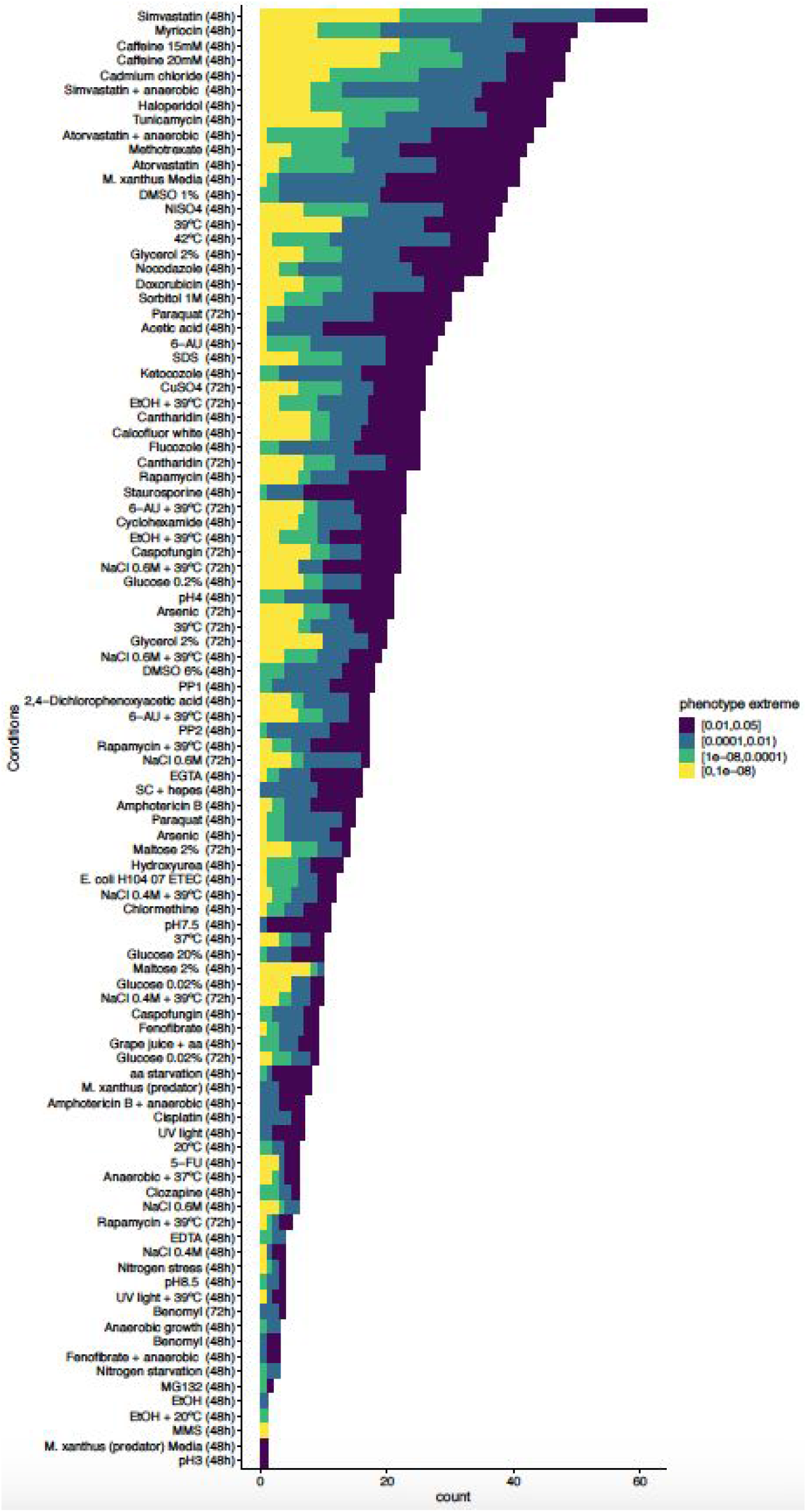
Number of mutant phenotypes per condition represented in different colors by their q-value

**Fig S4.**
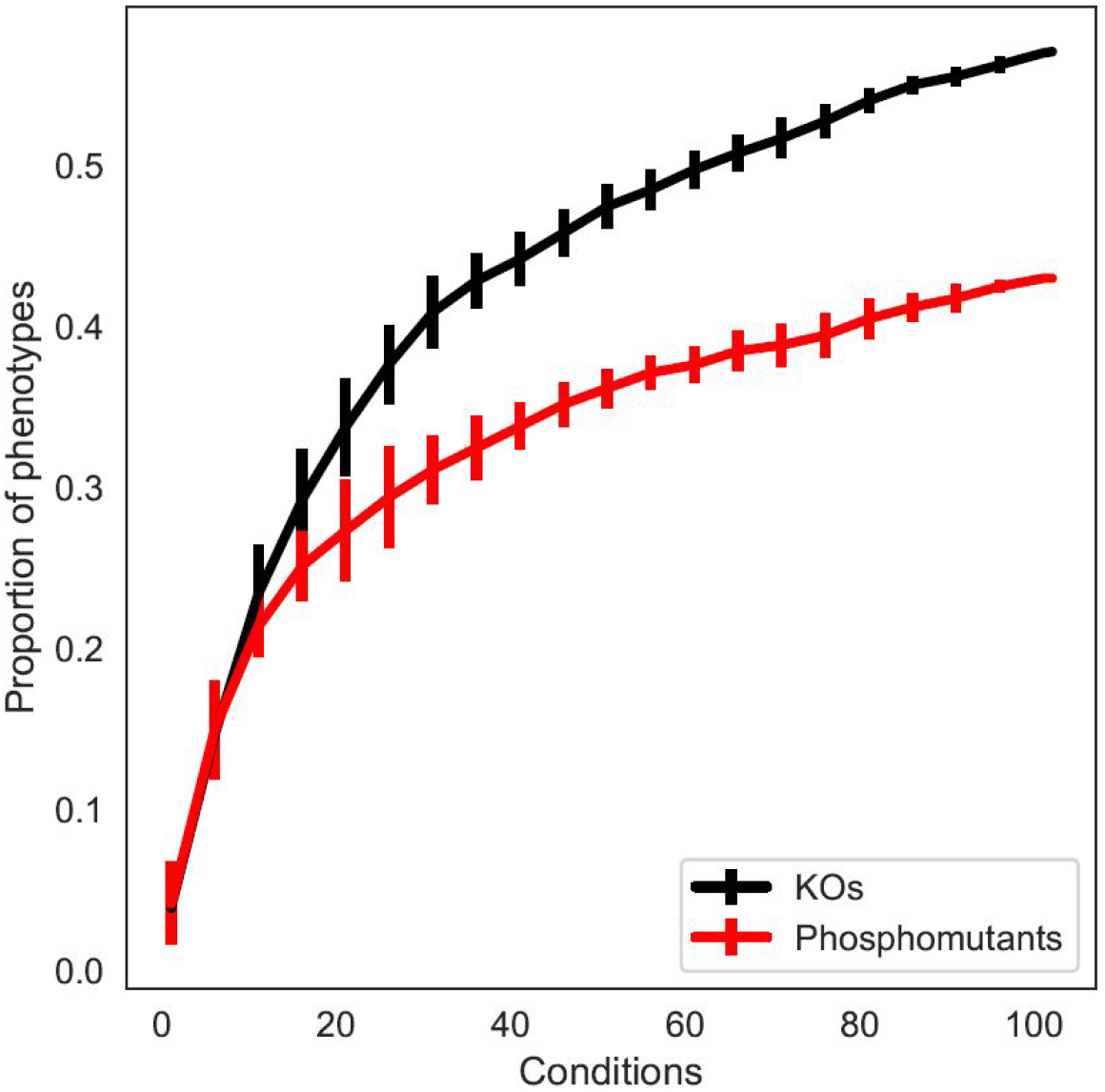
Rarefaction curves of the phenotypic screen for KOs and phospho-mutants. Proportion of phospho-mutants (red) and knockout strains (black) with phenotypes across conditions, plot shows both have not yet reached saturation.

**Fig S5.**
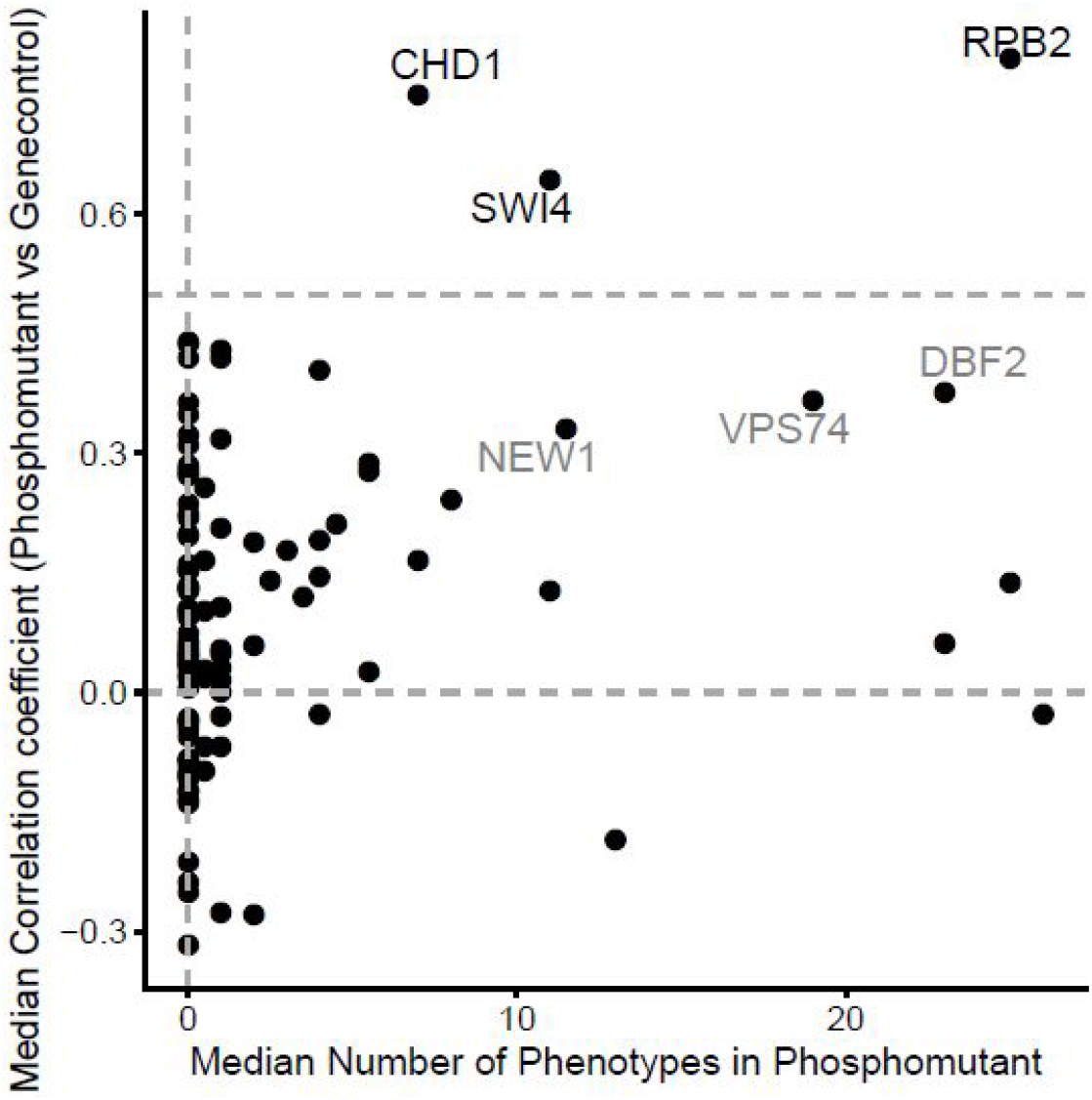
Phospho-mutant phenotypes vs median scar correlation phenotypes. Dashed line highlighting the genes *CHD1, SWI4* and *RPB2*), which show a correlations above the cutoff (r=0.5), and to a lesser extent *NEW1, VPS74* and *DBF2*, indicating the 18bp scar may have an impact on the observed phenotypes.

**Fig S6.**
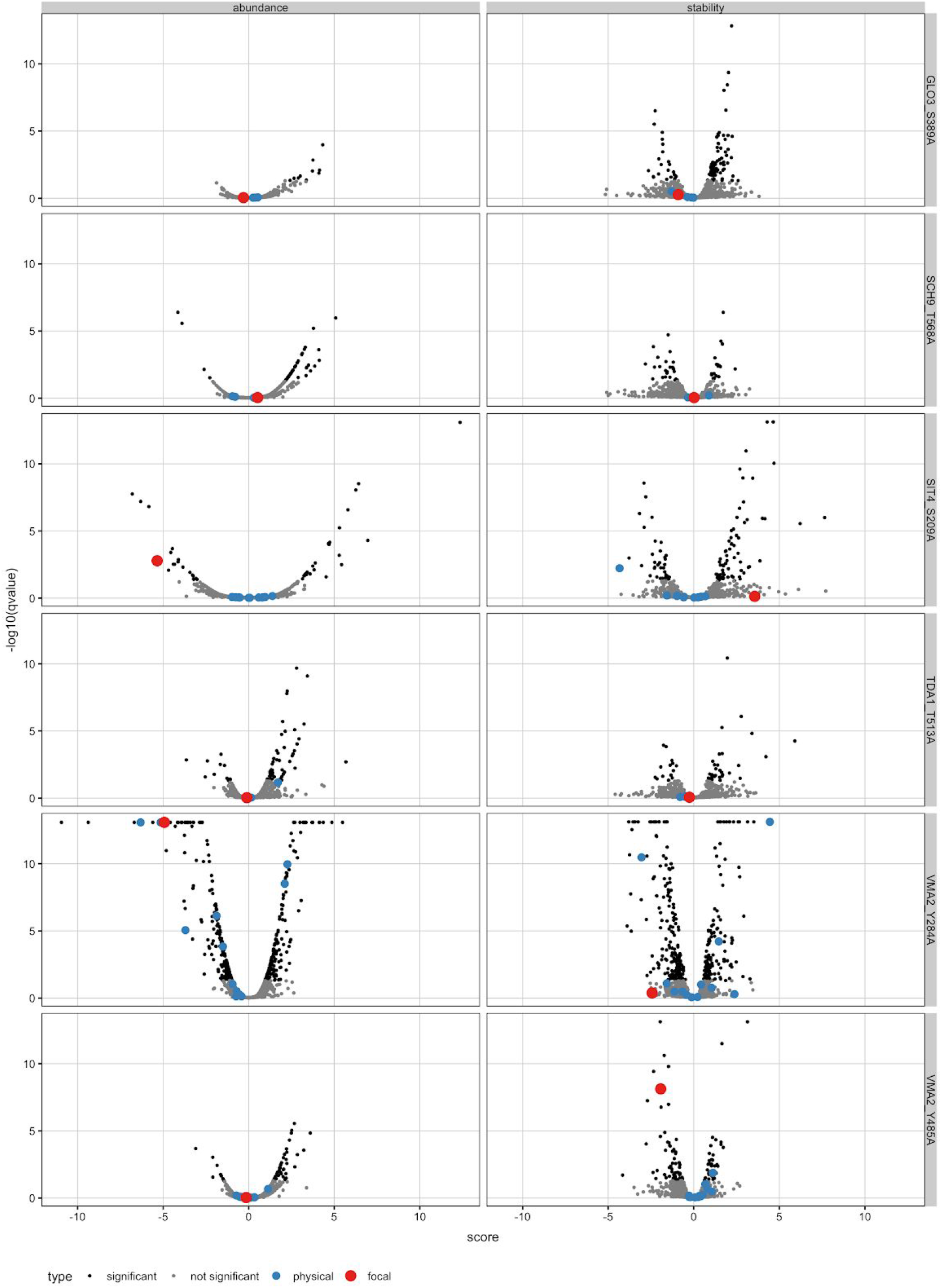
Volcano plots from TPP experiments highlighting protein interacting partners

**Fig S7.**
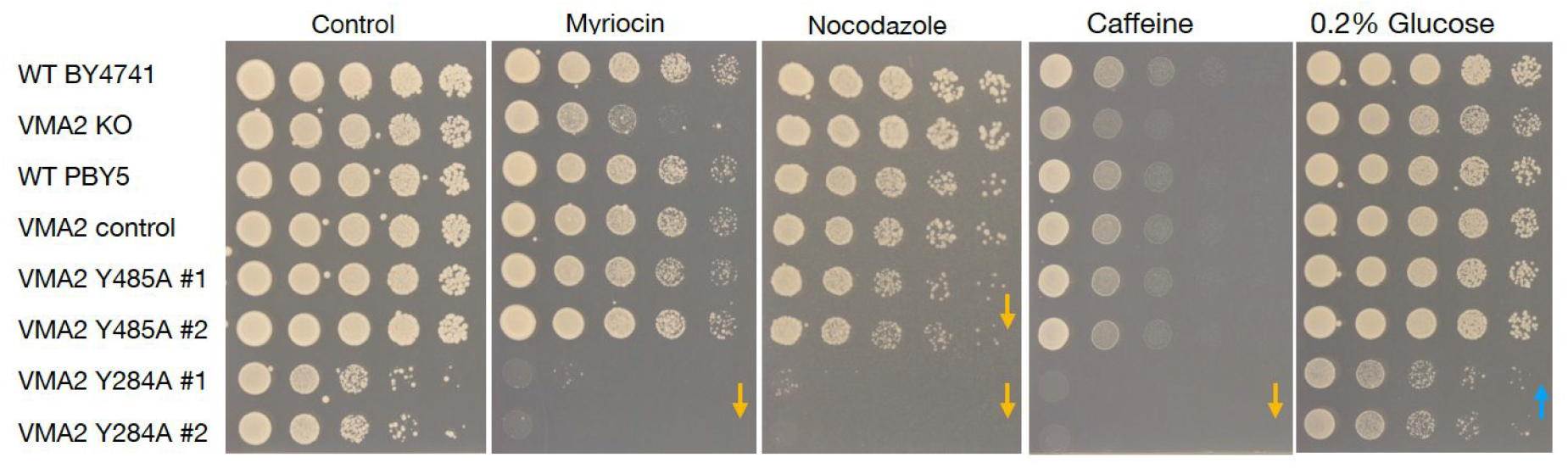
Phenotype validation of VMA2 strains by spot assays

**Fig S8.**
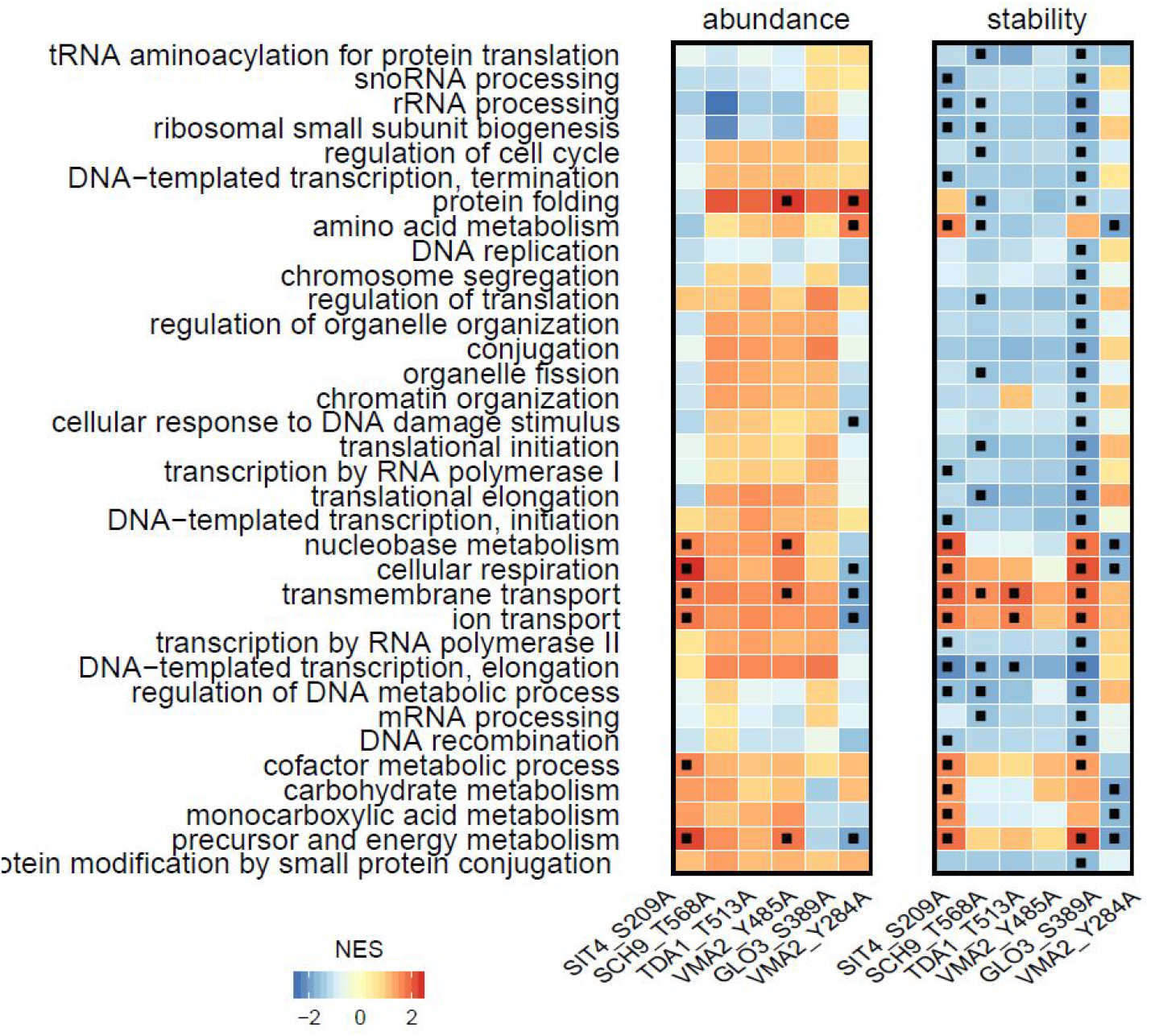
GO enrichment for the subset of phospho-mutants tested by TPP

**Fig S9.**
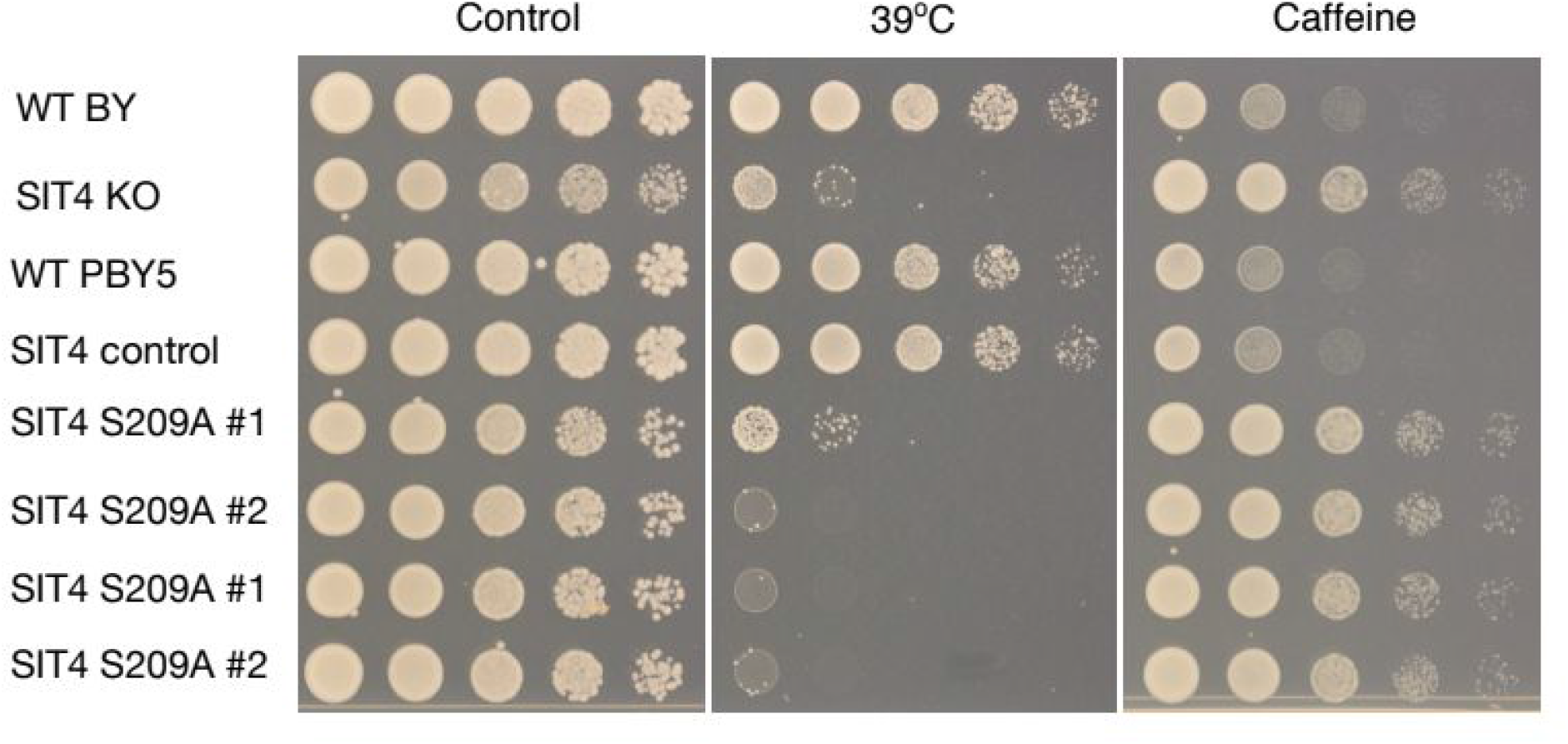
Phenotype validation of SIT4 strains by spot assays

## Acknowledgements

Nevan Krogan for generously providing strains. Michael Knop for generously providing plasmids. Natalia Gabrielli for providing nitrogen stress media. Magdalena Gierlach and Nadja Nepke from the EMBL Lab kitchen for their help making media and pouring screening plates. Rose Gathungu and Prasad Phapale from the EMBL Metabolomics Core Facility. Kevin Roy and Judit Villen for critical reading of the manuscript. This study has been funded by EMBL core funding and a Starting Grant Award from the European Research Council (ERC-2014-STG 638884 PhosFunc).

